## Supplemental Materials for "Within-species floral evolution reveals convergence in adaptive walks during incipient pollinator shift"

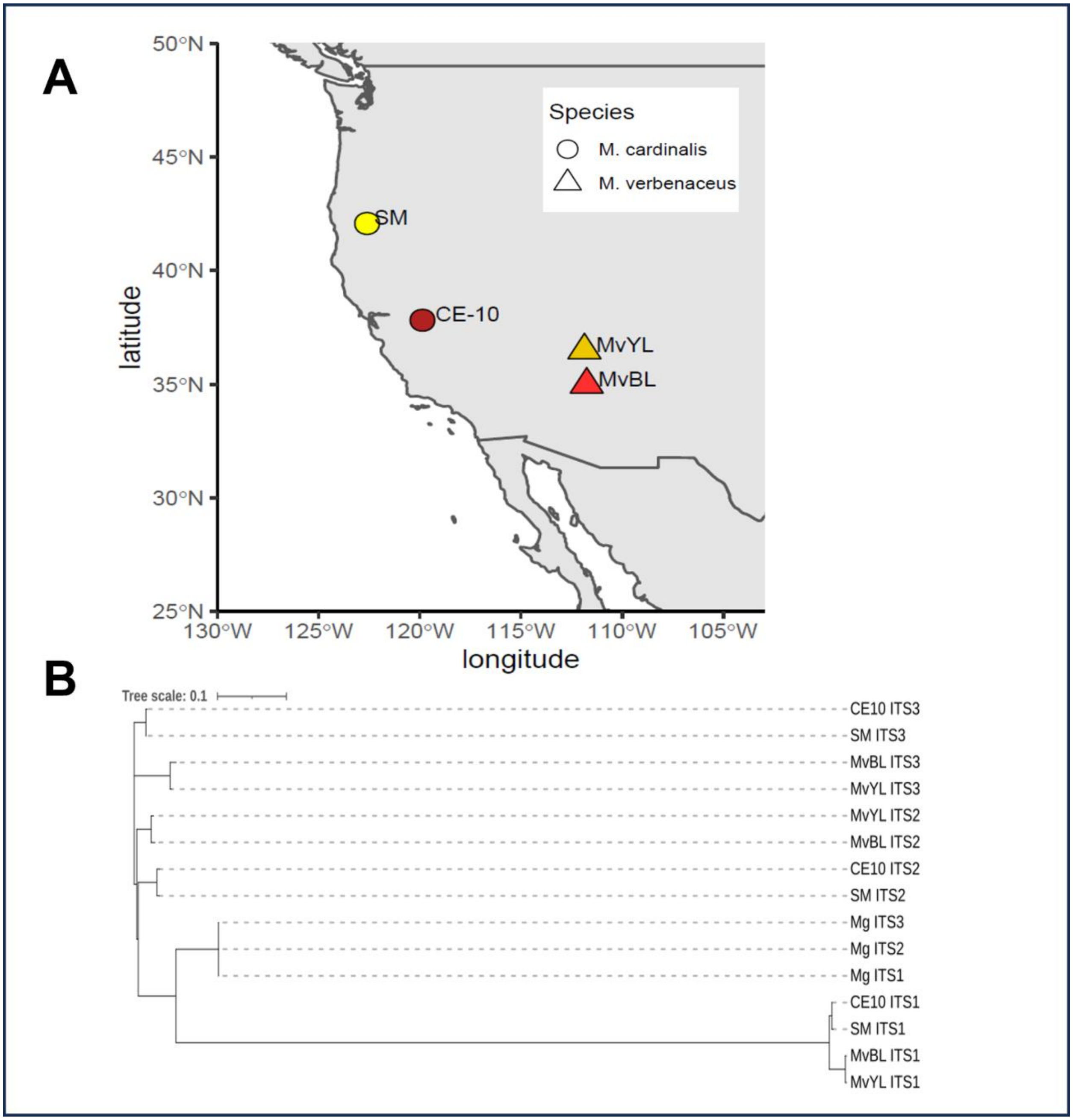


**Figure S1. (A)** Map of approximate collection locations of seed of focal lines, showing geographically distant populations of novel yellow morphs, consistent with their repeated independent origins. **(B)** Evolutionary relationships of focal lines of *M. verbenaceus* (MvBL and MvYL) and *M. cardinalis* (CE-10 and SM), with congener *M. guttatus* (Mg) as outgroup; the evolutionary history was inferred by using the Maximum Likelihood method and Tamura-Nei model on 3 ITS sequences from the genomes of the four focal morphs and outgroup *Mimulus*

*guttatus* (Mg). Analysis confirms that yellow morphs (MvYL and SM) are most closely related to their red conspecific lines (MvBL and CE-10, respectively), again consistent with their repeated independent origins. Abbreviations: MvYL = MvY, MvBL = MvR, SM = McR, CE-10 = McR.

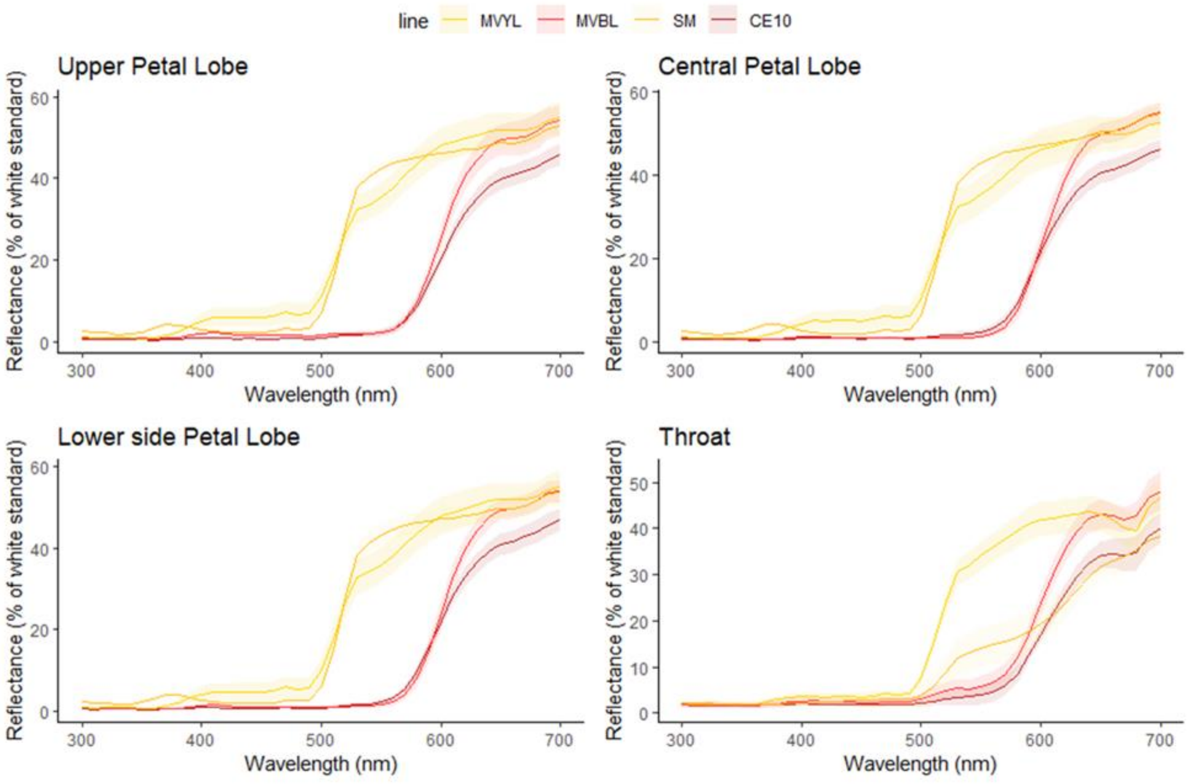


**(A)**

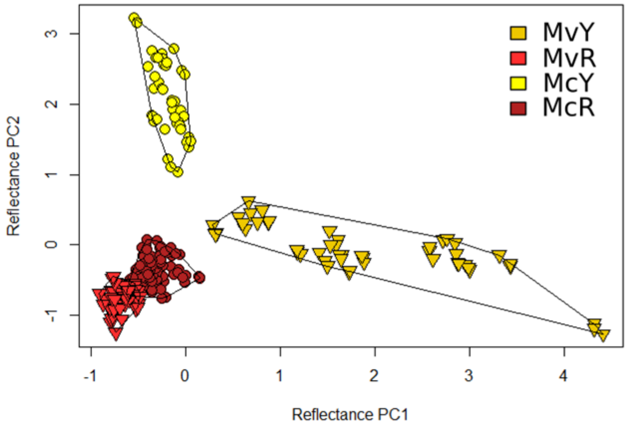


**(B)**

Figure S2. (A) Reflectance curves of all four corolla tissues measured. (B) Plot of PC1 and PC2 of PCA from reflectance data of central petal lobes of all plant lines. Abbreviations: MvYL = MvY, MvBL = MvR, SM = McR, CE-10 = McR.

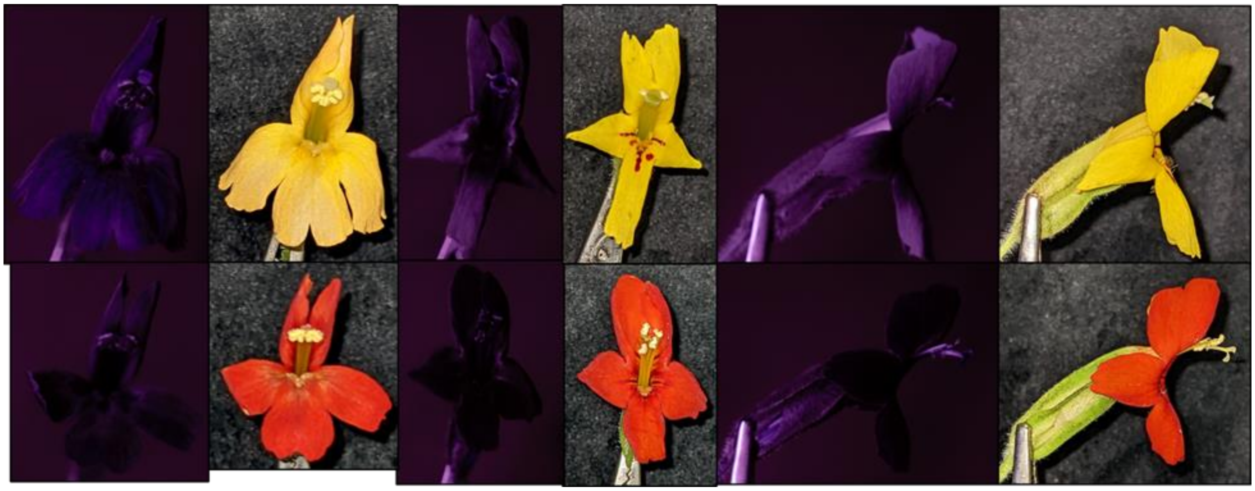

Figure S3. UV photography alongside visual spectrum photography of the same flowers from each plant line. Left two panes: *M. verbenaceus* (MvY, left top row; MvR, left bottom row). Lines of *M. cardinalis* (McY, center and right top row; McR, center and right bottom row) are also shown in side-view given their highly reflexed petal lobes.

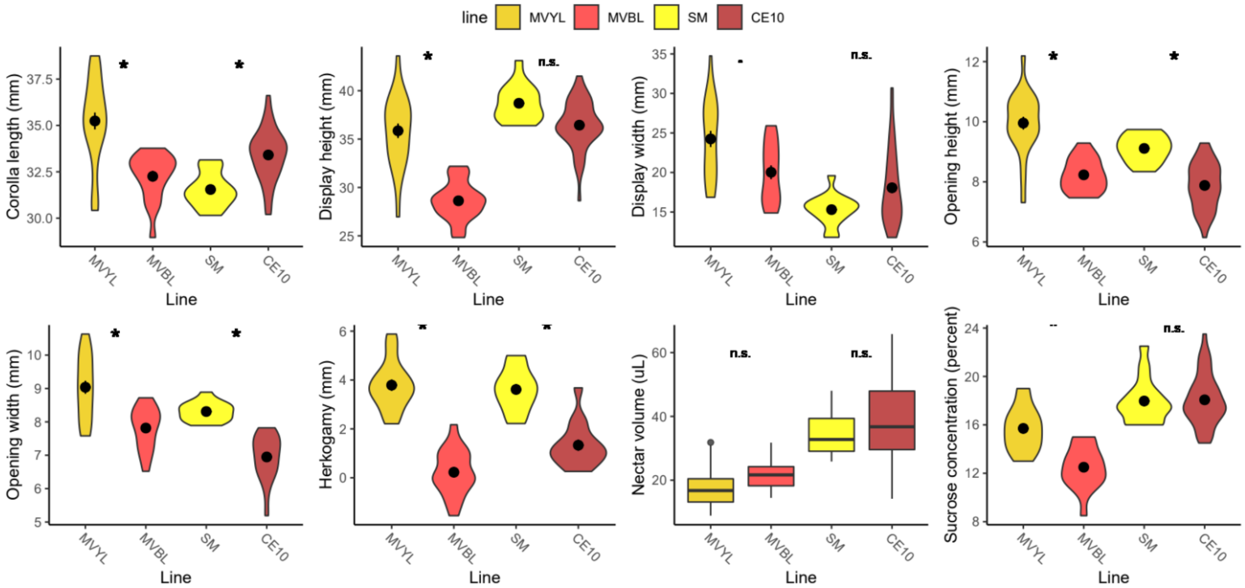

Figure S4. Violin plots of all measured floral morphology and nectar traits by plant line. Dot shows mean and bars +/- standard error. Abbreviations: MvYL = MvY, MvBL = MvR, SM = McR, CE-10 = McR.

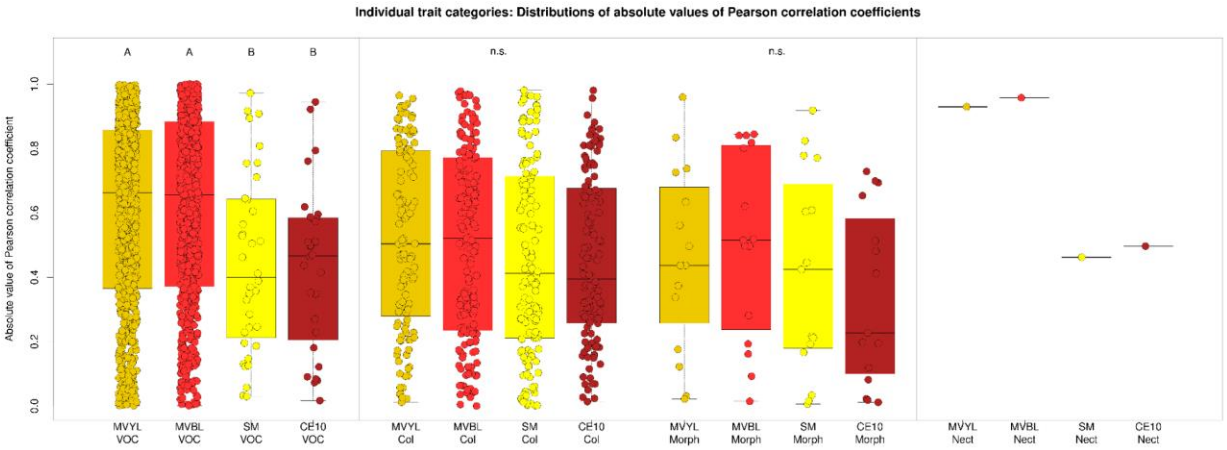


Figure S5. Distributions of absolute Pearson Correlation Coefficient values within trait categories (scent, color, morphology, nectar) sorted by trait type. VOC: volatile organic compounds; Col: pigment and reflectance spectrophotometry (latter values from principal component analysis); Morph: morphological traits; Nect: nectar traits. Statistical analysis of Nec was not performed given only one correlation value was available per line. Letters above or below boxes indicate statistically significant differences. n.s., not significant. Abbreviations: MvYL = MvY, MvBL = MvR, SM = McR, CE-10 = McR.

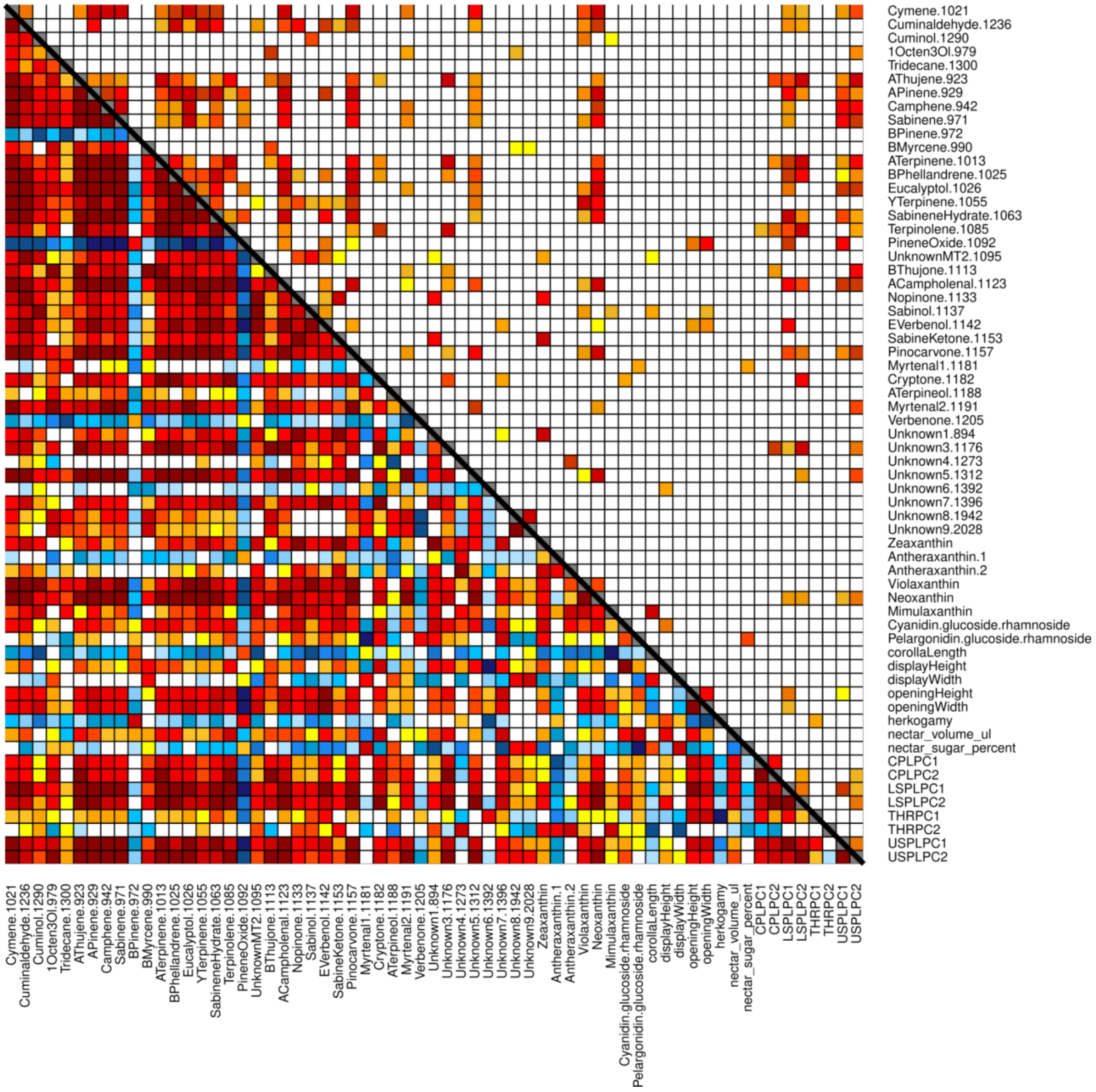


Figure S6. All-trait correlations in MvY. The lower left triangle displays Pearson correlation coefficients (blue = negatively correlated, yellow to red = positively correlated, darker colors are more strongly correlated) and the upper right triangle displays the p-values of correlations (white is $p > 0.05$, darker red is more significant). CPLPC1-2, central petal lobe reflectance PC1-2; LSPLPC1-2, lower side petal lobe reflectance PC1-2; THRPC1-2, corolla throat reflectance PC1-2; USPLPC1-2, upper side petal lobe reflectance PC1-2.

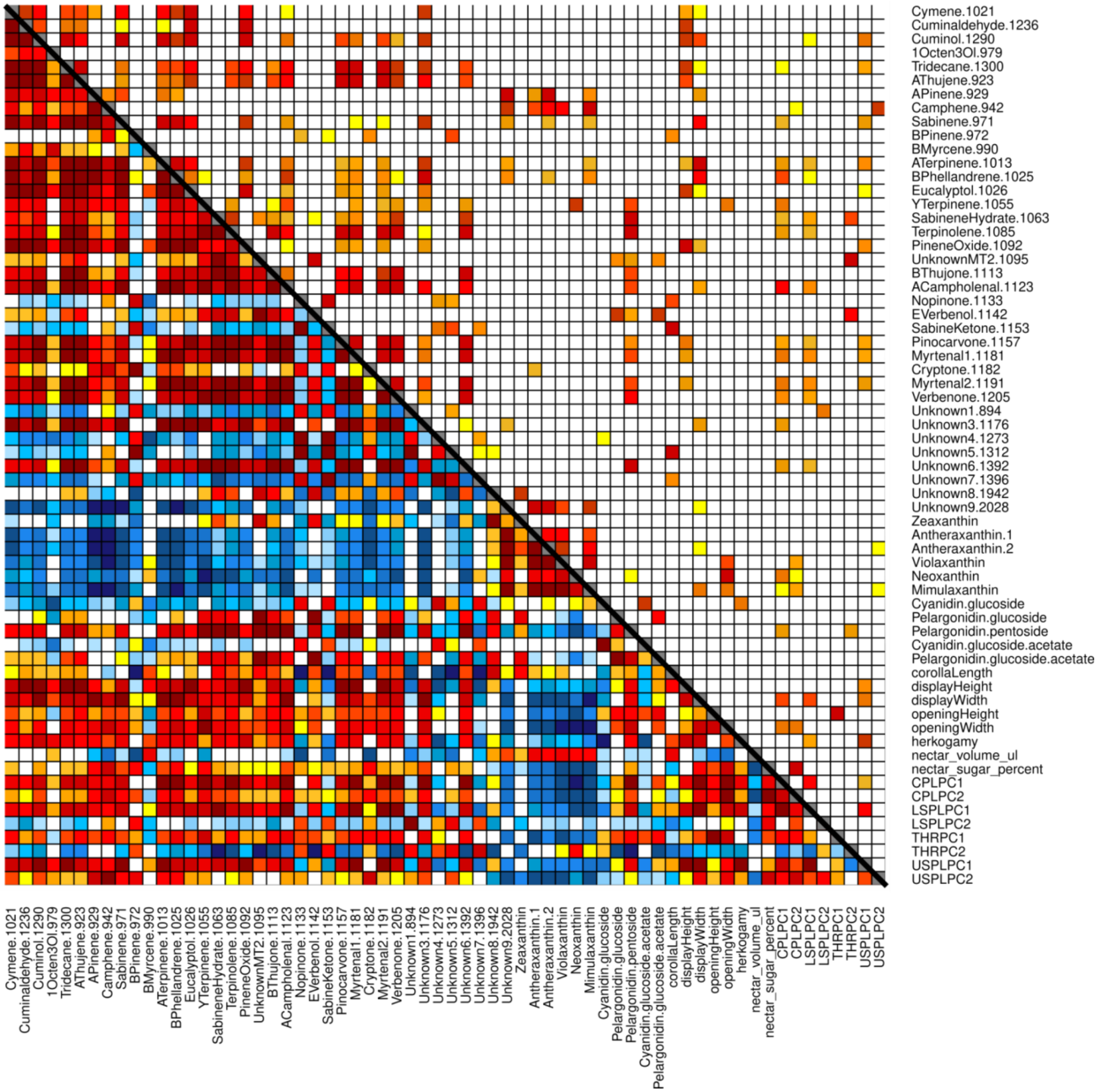


Figure S7. All-trait correlations in MvR. The lower left triangle displays Pearson correlation coefficients (blue = negatively correlated, yellow to red = positively correlated, darker colors are more strongly correlated) and the upper right triangle displays the p-values of correlations (white is $p > 0.05$, darker red is more significant). CPLPC1-2, central petal lobe reflectance PC1-2; LSPLPC1-2, lower side petal lobe reflectance PC1-2; THRPC1-2, corolla throat reflectance PC1-2; USPLPC1-2, upper side petal lobe reflectance PC1-2.

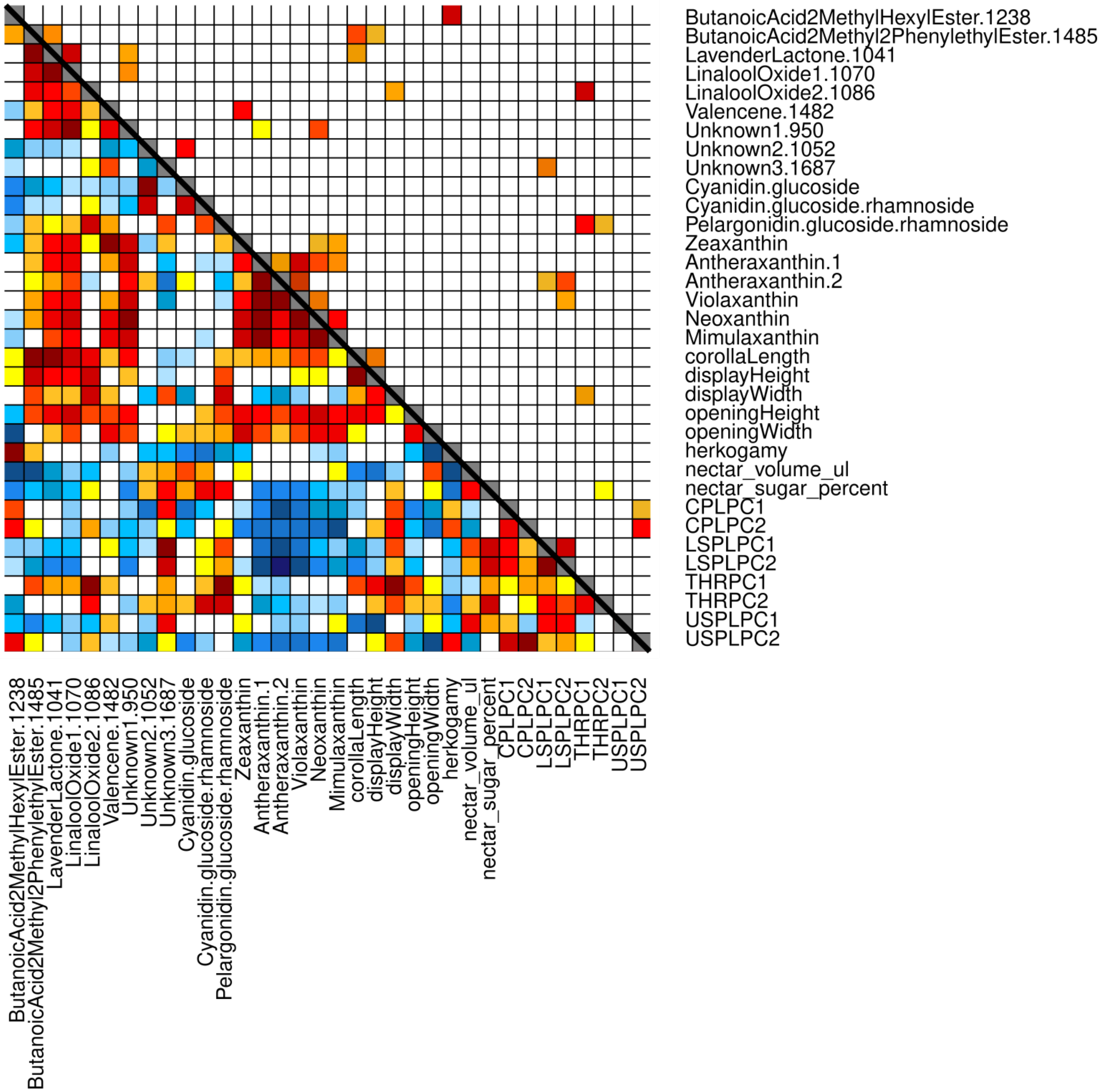


Figure S8. All-trait correlations in McY. The lower left triangle displays Pearson correlation coefficients (blue = negatively correlated, yellow to red = positively correlated, darker colors are more strongly correlated) and the upper right triangle displays the p-values of correlations (white is $p > 0.05$, darker red is more significant). CPLPC1-2, central petal lobe reflectance PC1-2; LSPLPC1-2, lower side petal lobe reflectance PC1-2; THRPC1-2, corolla throat reflectance PC1-2; USPLPC1-2, upper side petal lobe reflectance PC1-2.

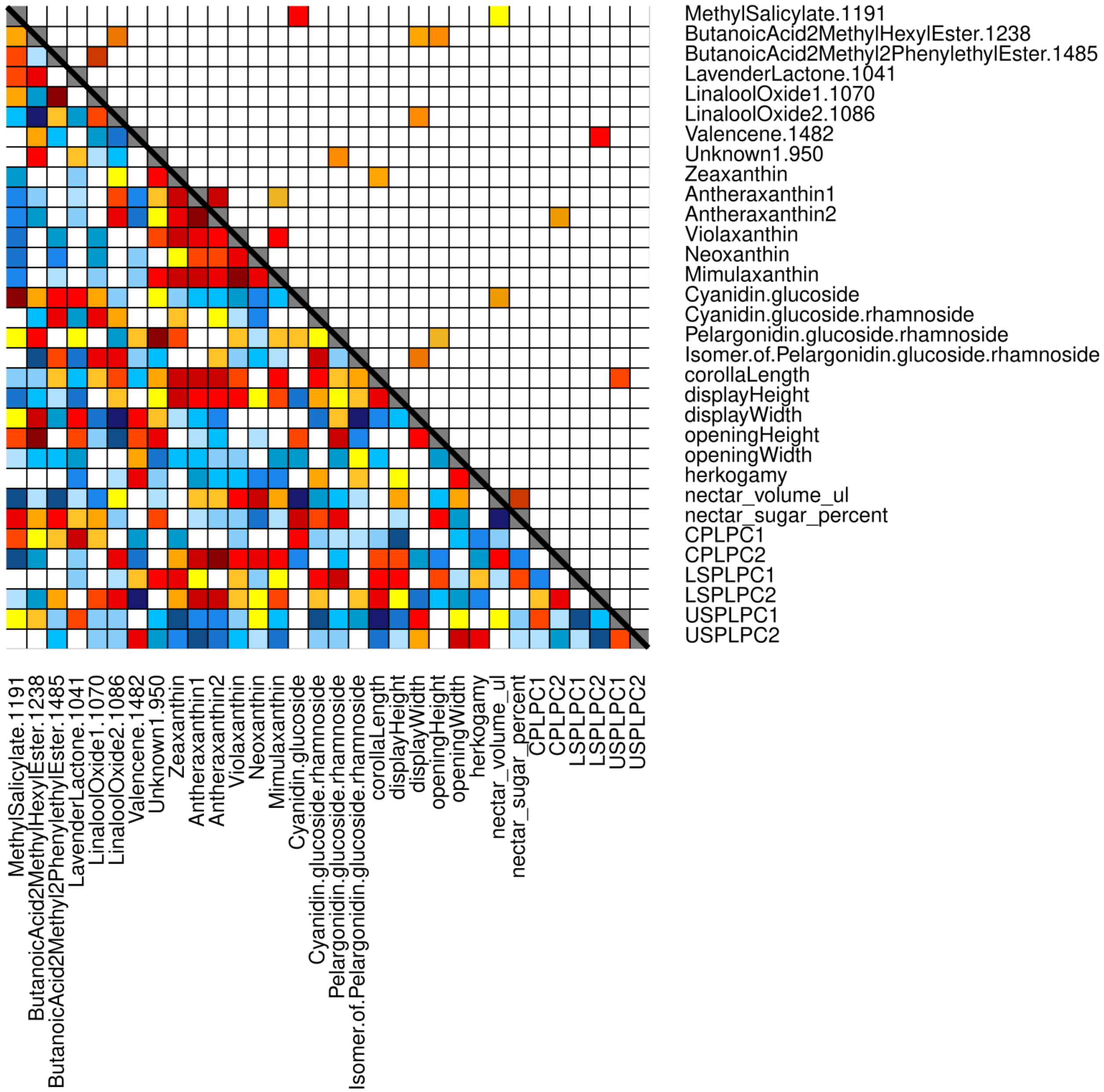


Figure S9. All-trait correlations in McR. The lower left triangle displays Pearson correlation coefficients (blue = negatively correlated, yellow to red = positively correlated, darker colors are more strongly correlated) and the upper right triangle displays the p-values of correlations (white is $p > 0.05$, darker red is more significant). CPLPC1-2, central petal lobe reflectance PC1-2; LSPLPC1-2, lower side petal lobe reflectance PC1-2; USPLPC1-2, upper side petal lobe reflectance PC1-2.

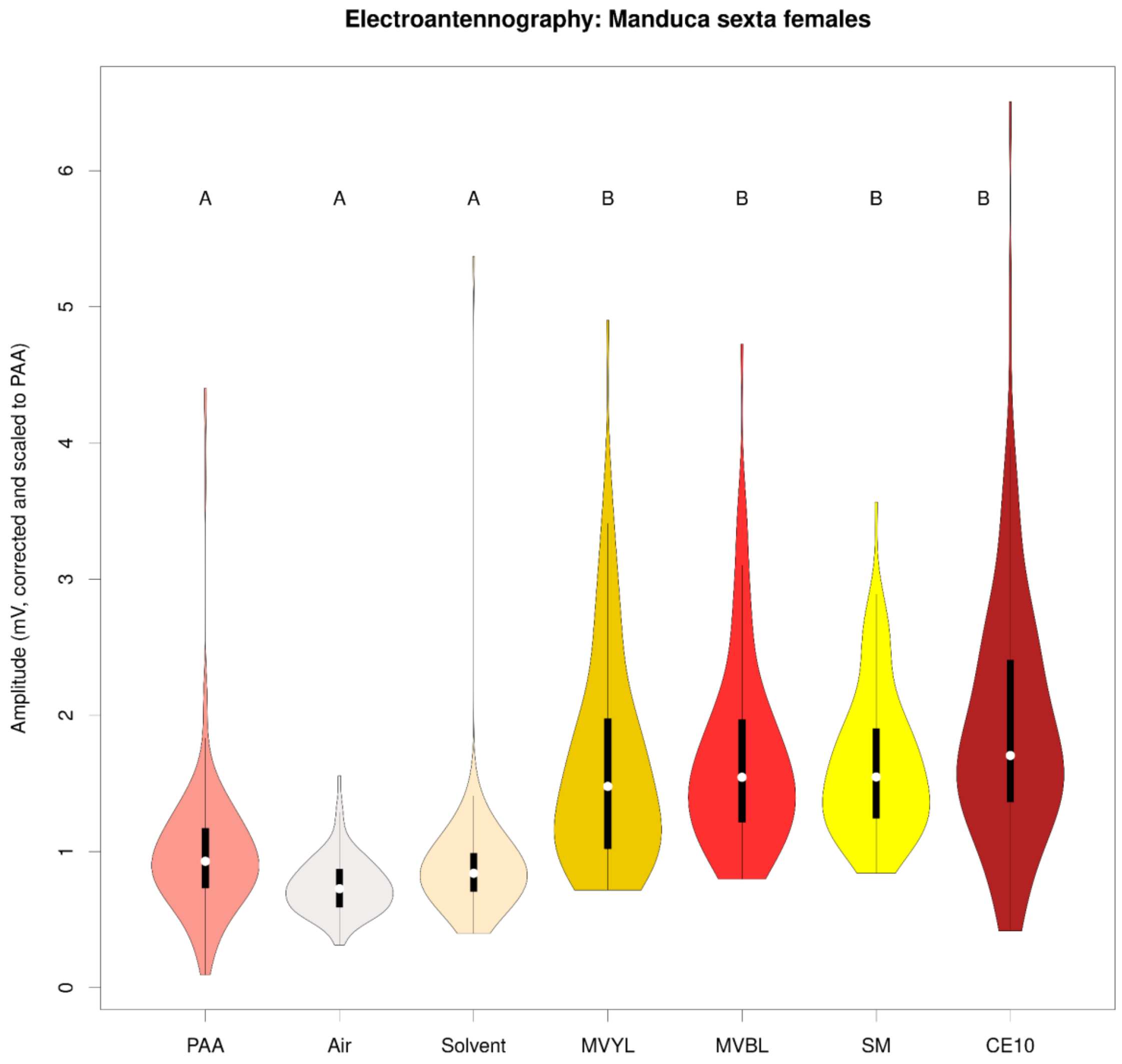


Figure S10: EAG responses to scent stimuli by seven female *Manduca sexta* hawkmoths, including phenylacetaldehyde (PAA, a floral VOC known to be detectable by *Manduca sexta*), air (negative control), extraction solvent alone, and floral scent extractions of MVYL, MVBL, SM, and CE10. Letters above violins indicate statistically significant differences. Abbreviations: MvYL = MvY, MvBL = MvR, SM = McR, CE-10 = McR.

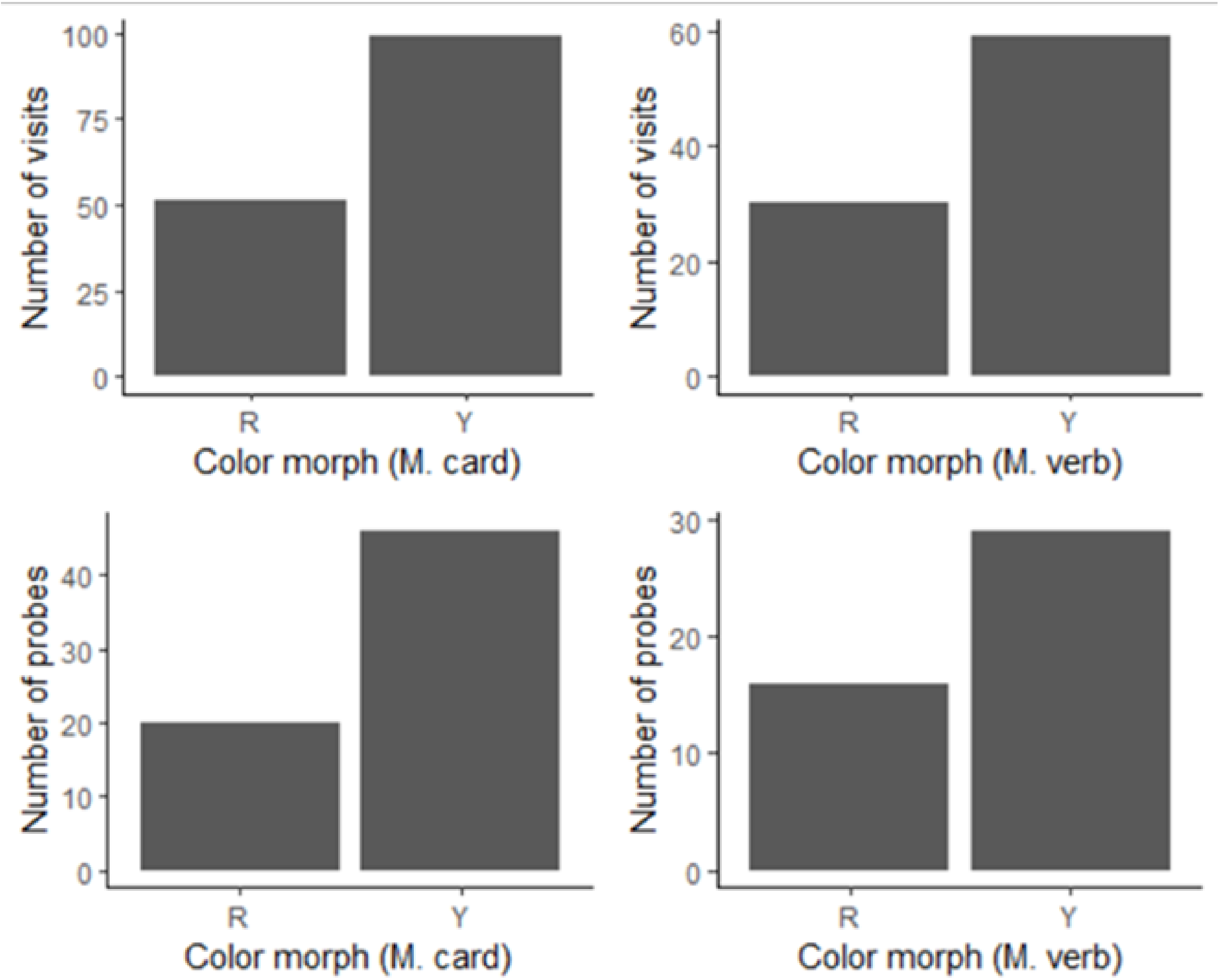


Figure S11. Total number of floral visits by bumblebees to each floral color morph for each within-species pairwise comparison, pooled across all bumblebees/trials.

Figure S12. Flavonoid Accumulation.

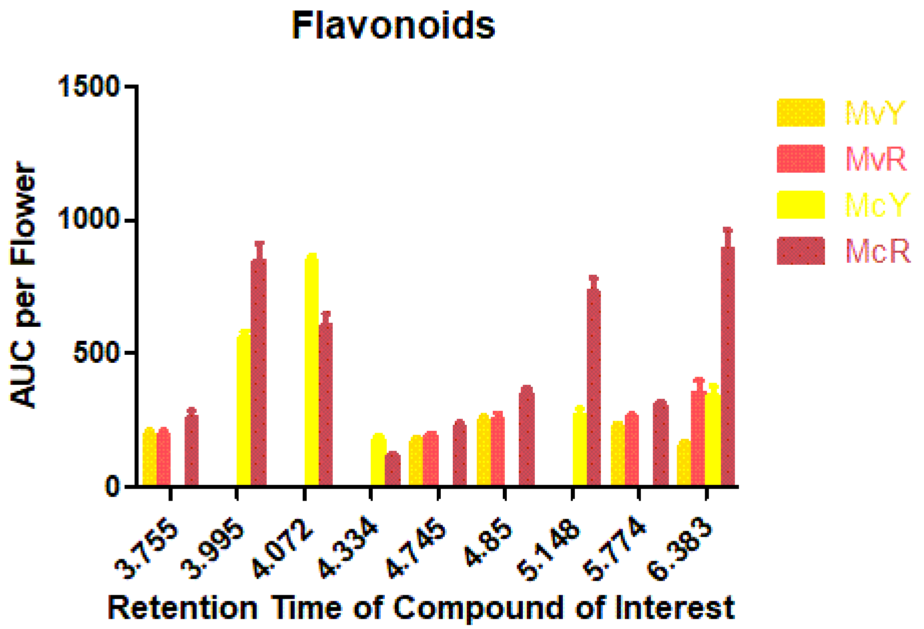


Figure S13. NMDS of whole transcriptomes of the four focal morphs visualized in Degust.

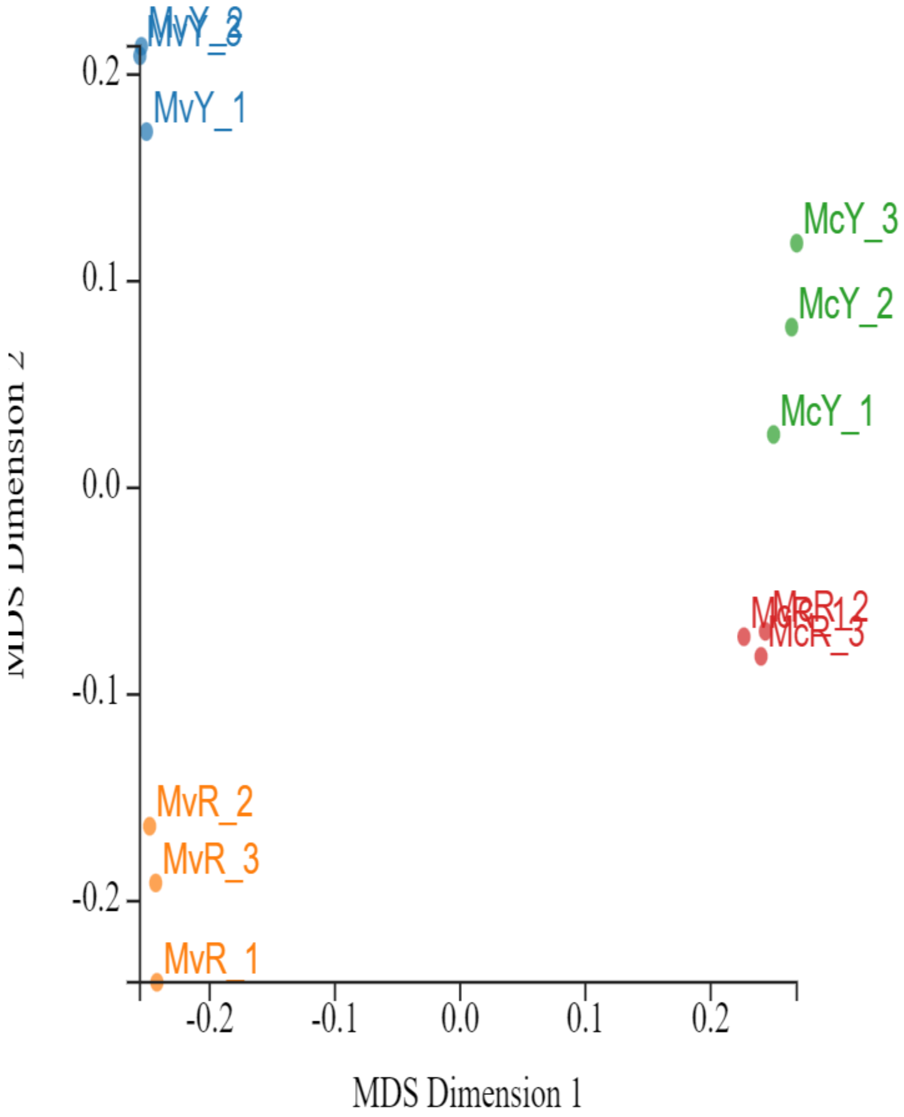


Figure S14. Gene expression data (mapped read counts) from six floral scent genes previously identified in *Mimulus* section *Erythranthe* with *Mimulus verbenaceus* MVBL gene ID listed. Letters above or below boxes indicate statistically significant differences. Abbreviations: MvYL = MvY, MvBL = MvR, SM = McR, CE-10 = McR.

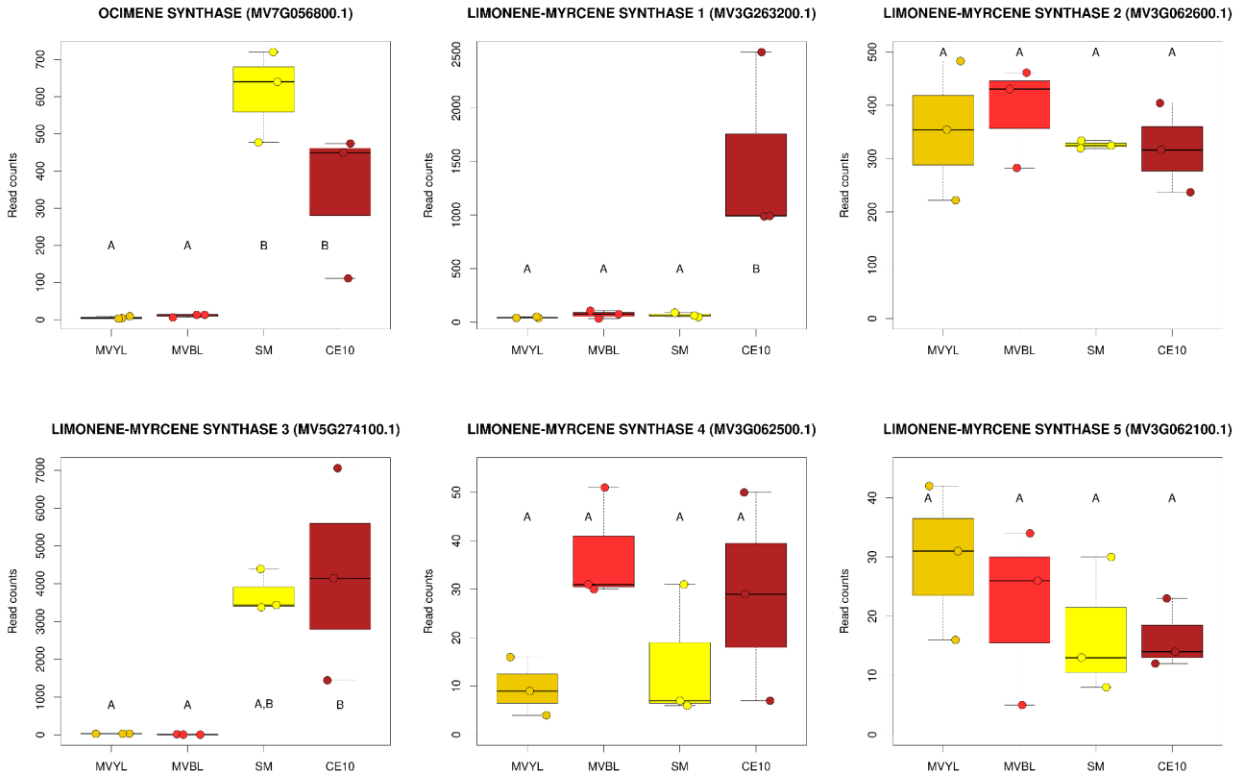


**SUPPLEMENTAL TABLES [please see separate Excel file for table contents]**

**TABLE S1.** Sample sizes and plant IDs used for each set of experiments.

**TABLE S2.** Absorbance for total anthocyanins (at 525nm) and total carotenoids (at 450nm) by line nested within species. Absorbance values were divided by average mass (g) of corolla tissues per line.

**TABLE S3.** Pairwise contrasts among floral morph/lines nested within species, based on linear mixed models with individual plant as a random effect. Floral morphology measurements in mm.

**TABLE S4.** Pairwise contrasts among *Bombus terrestris* EAG responses to scent stimuli of all four floral morph lines (MVYL, MVBL, SM, CE10), negative control (air), extraction solvent, and phenylacetylaldehyde (PAA, concentration 0.5 ng/uL), known to elicit a response in *Bombus terrestris ssp. audax*. All response values are corrected and scaled to PAA.

**TABLE S5.** Pairwise contrasts among *Manduca sexta* EAG responses to scent stimuli of all four floral morph lines (MVYL, MVBL, SM, CE10), negative control (air), extraction solvent, and phenylacetylaldehyde (PAA, concentration 0.5 ng/uL), known to elicit a response in *Manduca sexta*.

**Table S6**. Two-way ANOVA of anthocyanin biosynthesis gene expression of all four floral morphs, generated in Graphpad Prism v5.04. Swift, M. L. (1997). GraphPad prism, data analysis, and scientific graphing. Journal of chemical information and computer sciences, 37(2), 411-412.

**Table S7**. Two-way ANOVA of anthocyanin biosynthesis regulators of all four floral morphs, generated in Graphpad Prism v5.04. Swift, M. L. (1997). GraphPad prism, data analysis, and scientific graphing. Journal of chemical information and computer sciences, 37(2), 411-412.

**Table S8**. Two-way ANOVA of individual flavonoids of all four floral morphs depicted as their retention times, generated in Graphpad Prism v5.04. Swift, M. L. (1997). GraphPad prism, data analysis, and scientific graphing. Journal of chemical information and computer sciences, 37(2), 411-412.

**Table S9**. One-way ANOVAs of individual carotenoids of all four floral morphs, generated in Graphpad Prism v5.04. Swift, M. L. (1997). GraphPad prism, data analysis, and scientific graphing. Journal of chemical information and computer sciences, 37(2), 411-412.

**Table S10.** Two-way ANOVA of carotenoid biosynthesis regulators of all four floral morphs, generated in Graphpad Prism v5.04. Swift, M. L. (1997). GraphPad prism, data analysis, and scientific graphing. Journal of chemical information and computer sciences, 37(2), 411-412.

**Table S11**. Statistical Analyses of genomic variants by 50kb window, per chromosome. Generated in Graphpad Prism v5.04. Swift, M. L. (1997). GraphPad prism, data analysis, and scientific graphing. Journal of chemical information and computer sciences, 37(2), 411-412.

**Table S12**. ITS Fasta sequences used to generate Maximum Likelihood Tree Figure 1E.
